# Systemic immune alterations in a murine experimental model of osteoarthritis

**DOI:** 10.1101/2025.11.22.689773

**Authors:** Célia Seillier, Sybille Brochard, Karim Boumediene, Denis Vivien, Brigitte Le Mauff, Olivier Toutirais, Catherine Baugé

## Abstract

Osteoarthritis (OA) is accompanied by an inflamed synovium containing macrophages, dendritic cells, T and B lymphocytes. Macrophages predominate and drive cytokine-mediated cartilage catabolism, while T cells and B cells, though fewer, may shape chronic adaptive responses. However, systemic immune contributions, particularly within peripheral lymphoid organs such as the spleen, remain poorly characterized. Our study aims to profile systemic immune changes in experimental OA induced by injection of mono-iodoacetate (MIA) in mouse paw. At day 56 post-OA induction, analysis of splenocytes showed that macrophages and conventional dendritic cells (cDC1 and cDC2) displayed a significant downregulation of MHCII expression, suggesting a negative feedback mechanism that limits chronic T cell activation. OA is also associated with an increase in total DCs including mainly MHCII negative tolerogenic DCs. Notably, while the proportion of CD11b^-^ tolerogenic DCs was reduced, CD11b^low^ tolerogenic DCs markedly expanded in OA animals. Expression level of the CD11b integrin was upregulated on macrophages and cDC2 in MIA-induced OA mice potentially facilitating their adhesion and migration toward inflamed joint. OA mice showed a significant reduction in total splenic leukocytes, primarily due to a loss of B cells, while total T cell numbers remained stable. However, T cell composition shifted: CD4^+^ T cells including activated and regulatory subsets decreased, whereas activated CD8^+^ T cells increased. This indicates a systemic imbalance favoring cytotoxic over regulatory immune activity, possibly linked to chronic immune stress or redistribution of lymphocytes to inflamed joint. In conclusion, our data reveals that chronic OA induces a coordinated remodeling of systemic innate and adaptive immunity. These systemic immune dysregulations could reveal new biomarkers or therapeutic targets.

## Introduction

Osteoarthritis (OA) is a leading cause of chronic musculoskeletal disability worldwide. It affects roughly 7-10% of the global population and is especially prevalent in older adults. OA patients suffer progressive joint pain, stiffness, and loss of function, driving large healthcare and socio-economic burdens [1]. Historically viewed as “wear-and-tear” degradation of cartilage, OA is now recognized to involve active inflammatory processes in multiple joint tissues [2–4]. Indeed, synovial inflammation (synovitis) accompanies even early-stage OA and predicts radiographic progression and pain severity [5]. Pathologically, OA involves not only cartilage loss and subchondral bone changes, but also synovial lining hyperplasia and immune cell infiltration [6]. Importantly, OA is increasingly characterized as a systemic low-grade inflammatory disease rather than a purely degenerative condition [4].

OA joints contain a mix of innate and adaptive immune cells [7–9]. Macrophages are the most abundant immune cells in osteoarthritic synovium. In OA, blood-derived monocytes infiltrate the joint and polarize into classically activated (M1) macrophages that secrete high levels of TNF, IL-1β, IL-6 and matrix metalloproteinases, driving cartilage catabolism. In contrast, alternatively-activated (M2) macrophages are fewer, and M1/M2 imbalances promote chronic synovial inflammation in OA [10]. Recent analyses of human OA synovium confirm marked increases in both M1 and M2 macrophages, as well as in activated dendritic cells (DCs) and mast cells, compared to healthy tissue [6,11]. These innate cells cooperate with complement, chemokines and inflammasome pathways to amplify joint inflammation [12,13]. DCs appear also in OA joints but are relatively rare compared to macrophages. DC-like cells have been identified in synovial fluid and synovium of early knee OA, though they do not form dense aggregates as in rheumatoid arthritis [11]. Nonetheless, activated DCs in OA tissue can present cartilage-derived antigens and sustain local inflammation [6,11].

T lymphocytes are readily detected in OA synovial tissue and fluid [14,15]. Analyses show that OA synovium contains significantly more CD3^+^ T cells than healthy tissue. Both CD4^+^ (helper) and CD8^+^ (cytotoxic) T cell subsets, including Th1, Th17, and regulatory T cells, infiltrate OA joints [16]. These T cells produce pro-inflammatory cytokines (e.g. IFN-γ, IL-17) that promote synovial inflammation and may influence subchondral bone remodeling [17]. In experimental models, deficiency of CD4^+^ T cells attenuates post-traumatic OA, indicating that adaptive immunity contributes to disease pathogenesis [18]. B lymphocytes and antibodies also play a role in OA, though their contributions are less well characterized. Small numbers of B cells and plasma cells are present in OA synovium [19]. Clonal B cell expansions and autoantibodies to cartilage antigens have been reported in OA patients, suggesting adaptive humoral responses are activated. However, B cells constitute only a minority (≈5%) of synovial leukocytes in OA, and their precise functions (e.g. antigen presentation vs. antibody production) remain under study [19].

Despite clear joint inflammation, it remains unclear how OA affects systemic immunity. Observational studies suggest OA patients often have elevated systemic inflammatory indices: for example, blood counts show higher neutrophil-lymphocyte ratios and activated circulating monocytes, and serum levels of C-reactive protein (CRP) and IL-6 correlate with knee OA severity. Peripheral monocytes from knee-OA patients display an activated phenotype with a higher expression of the chemokine CCR2, major histocompatibility complex (MHC) class II molecules (HLA-DR) and the IgG Fc receptor (CD16). They also produce more TNF and IL-1β upon stimulation than controls [20]. Systemic biomarkers like the systemic immune-inflammation index have been linked to OA presence in large population cohorts, consistent with a low-grade inflammatory state beyond the joint. However, data regarding the activation status of immune cells in secondary lymphoid organs in OA are lacking.

In particular, spleen and lymph nodes (LNs), which are the central hubs of immune activity, have not been systematically studied in OA models or patients. In autoimmune arthritis (e.g. rheumatoid arthritis), draining LNs and spleen enlarge with expanded populations of activated T and B cells [21,22]. If similar changes might occur in OA, it has not been investigated yet. The gap in knowledge is significant: it is unknown whether chronic OA drives expansion or functional shifts in splenic or LN immune cell populations, and whether such systemic alterations influence or reflect joint pathology. Understanding systemic immunity in OA could reveal new biomarkers or therapeutic targets.

Given these considerations, we designed this study to profile systemic immune changes in an experimental murine OA model. Specifically, we focused on cells present in secondary lymphoid organs (spleen) in monosodium iodoacetate (MIA) induced OA mice.

## Material and methods

### Animals

Animal experiments were conducted in strict accordance with European Directive 2010/63/EU and French regulations (Decree 87/848) at the GIP CYCERON/CURB facility (Centre Universitaire de Ressources Biologiques, accreditation #D14118001). All procedures received prior authorization from the French Ministry of Education and Research as well as approval from the regional ethics committee CENOMEXA (APAFIS authorization #16185). Experimental procedures complied with the ARRIVE guidelines (Animal Research: Reporting of In Vivo Experiments; https://www.nc3rs.org.uk). Handling of animals followed internationally recognized ethical standards for laboratory animal care. Particular attention was given to the principles of the 3Rs (Refine, Reduce, Replace), ensuring that experiments were optimized, animal use was minimized whenever possible, and researchers were fully aware of their ethical responsibilities.

A total of 7 male C57BL/6J mice (8 weeks old, Janvier Labs, France) were used (n=3 sham mice and n=4 MIA induced mice). Animals were housed under controlled environmental conditions (temperature 23 ± 2 °C; 12-h light/dark cycle, reversed). Food and water were provided ad libitum, and cage bedding was replaced weekly. Experimental procedures were carried out between 08:00 and 17:00 in a room maintained under low light intensity (6 lx).

### OA induction

Osteoarthritis was induced by an intra-articular injection of 0.75 mg monosodium iodoacetate (MIA, Sigma-Aldrich) diluted in 10 μL of sterile physiological saline. Prior to the procedure, the working surface was disinfected with alcohol and a sterile field was prepared. Animals were anesthetized with 5% isoflurane in a mixture of 70% N_2_O/30% O_2_ for induction, followed by maintenance at 3% isoflurane under the same gas mixture. The total duration of anesthesia averaged 10 minutes, including both induction and recovery phases.

Mice were positioned in dorsal recumbency, and the right knee was disinfected with 2% alcoholic chlorhexidine (Gilbert, Hérouville-Saint-Clair, France). The joint was flexed to a 45° angle, and the articular space was located by visualizing the patellar tendon through the skin. A 10 μL injection was then administered over a 1-minute period directly into the synovial capsule using a 30G needle (Microlance 3, 30 G ½″, 0.3 × 13 mm; BD, Rungis, France) attached to a 25 μL Hamilton syringe (Giarmata, Romania). After needle withdrawal, the joint was gently mobilized to facilitate distribution of the solution. The injection site was subsequently rinsed with physiological saline to remove any residual MIA and prevent skin irritation.

At the end of the procedure, anesthesia was discontinued, and the animals were held manually until full recovery in order to prevent hypothermia. Their health status was monitored for one-hour post-injection, during which no signs of discomfort were observed.

At day 56, mice were euthanized by cervical dislocation after deep anesthesia (5% isoflurane, 70% N_2_O / 30% O_2_).

### Flow cytometry

To assess systemic inflammatory processes, spleens were collected and splenocytes dissociated and resuspended in 50 μL of staining buffer. Fc receptors were blocked with 10 μg/mL anti-CD16/CD32 antibodies (BD Biosciences) for 15 min at 4 °C. Cells were then labelled for cell surface markers with fluorochrome-conjugated monoclonal antibodies for 10 min in the dark at 4 °C. When necessary, intracellular staining was performed using an “inside stain” kit (Miltenyi Biotec). Prior to tube acquisitions, 7-AAD (420,404, BioLegend) was added 15 min before analysis by flow cytometry. The used antibodies were: CD16/32 (2.4G2, 553,142, BD Biosciences), CD11b (REA592 VioBlue, 130-113-810, Miltenyi Biotec), CD11c (REA754 APC, 130-110-839, Miltenyi Biotec), F4/80 (REA126 PE-Vio770, 130-118-459, Miltenyi Biotec), CD80 (REA983 APC-Vio770, 130-116-463, Miltenyi Biotec), CD86 (REA1190 PE, 130-122-129, Miltenyi), MHCII (REA813 FITC, 130-112-386, Miltenyi Biotec), CD38 (REA616 PerCP-Vio700, 130-109-260, Miltenyi Biotec), iNOS (REA982 APC, 130-116-423, Miltenyi Biotec), CD206 (C068C2 APC, 141,708, BioLegend), Arg-1 (A1EXF5 PE, 12-3697-82, Invitrogen), CD3e (145-2C11 BV510, 563024, BD Biosciences), CD45 (30-F11 PE-Cy7, 561868, BD Biosciences), CD45R/B220 (RA3-6B2 APC, 553092, BD Biosciences), CD4 (RM4-5 APC, 55305, BD Biosciences), CD8a (53-6.7 PE-Cy7, 552877, BD Biosciences), CD25 (7D4 BV421, 564571, BD Biosciences), CD44 (IM7 FITC, 561859, BD Biosciences), CD69 (H1.2F3 PE, 561932, BD Biosciences). Samples were analyzed on a FACSVerse (BD Biosciences) with FlowJo 7.6.5 (TreeStar Inc.).

## Results

### Phenotype of splenic macrophages in the MIA-induced OA model

We induced OA in mice by intra-articular injection of MIA. This well-established model in our laboratory resulted in severe OA, characterized by joint degradation and synovitis [23,24]. Then, we evaluated the impact of MIA-induced OA on systemic inflammation. Using flow cytometry analysis, we have characterized the phenotype of macrophages (F4/80^+^ CD11b^+^) in spleen of mice treated with MIA at day 56 (Fig 1A). The cell count and frequency of whole splenic macrophages were not affected by MIA treatment (Fig 1B, C). A clear decrease of MHCII expression associated with a low increase of CD86 expression were observed on whole MIA macrophages (MHCII median fluorescence intensity (MFI) Sham 7790 [6607-7860] *vs* MIA 4655 [4010-5963], *p=*0.0286; CD86 Sham 718 [679-743] *vs* MIA 784 [767-831], *p=*0.0286) (Fig 1C). In the MIA condition, the proportions of both CD80^+^ CD86^+^ and CD80^-^ CD86^+^ macrophages were increased among MHCII^-^ and MHCII^+^ subsets (Fig. 1D, F). While CD86 expression showed a slight increase on MHCII^-^ macrophages, no significant difference was observed on MHCII^+^ macrophages (Fig. 1E, G). Finally, there was a tendency for the CD11b marker to increase only on MHCII^+^ macrophages after MIA treatment (Fig 1G).

**Figure 1.**
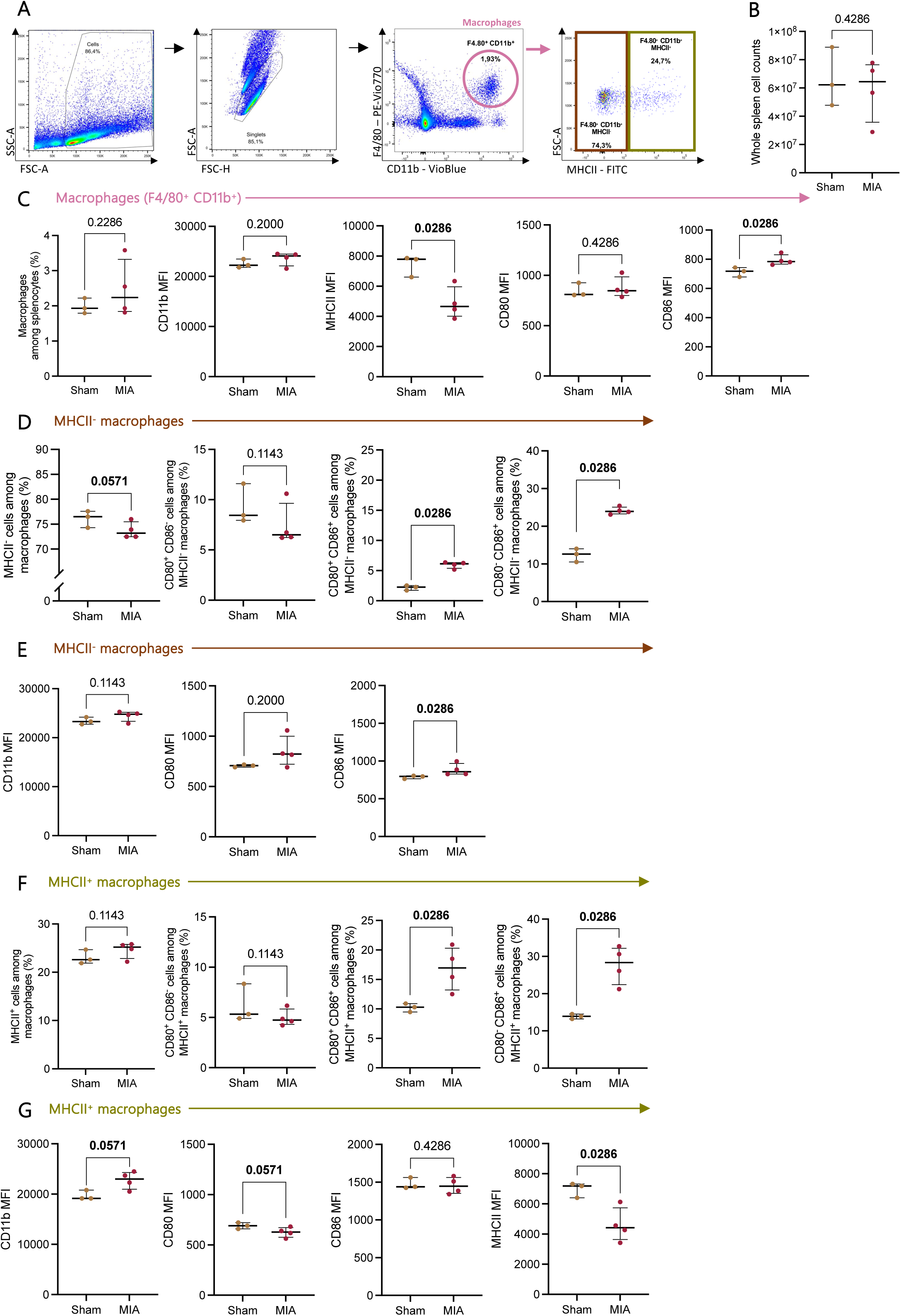
Modulation of macrophage activation status in the MIA-induced OA model. (**A**) Representative gating strategy for splenic macrophages (F4/80⁺ CD11b⁺), and their subsets according to MHCII expression (MHCII⁺ and MHCII⁻ macrophages). (**B**) Total spleen cell counts in MIA and sham mice. (**C**) Quantification of total macrophages among splenocytes (%) and analysis of surface marker expression (median fluorescence intensity; MFI - CD11b, MHCII, CD80, CD86). (**D, E**) Characterization of MHCII^-^ macrophages. Quantification of MHCII^-^, CD80^+^ CD86^+^ and CD80^-^ CD86^+^ macrophages among splenocytes (%) and analysis of surface marker expression (MFI - CD11b, MHCII, CD80, CD86). (**F, G**) Characterization of MHCII^+^ macrophages. Quantification of MHCII^+^, CD80^+^ CD86^+^ and CD80^-^ CD86^+^ macrophages among splenocytes (%) and analysis of surface marker expression (MFI - CD11b, MHCII, CD80, CD86). Data are shown as individual values with median [25% percentile-75% percentile] (Sham n=3; MIA n=4; Mann-Whitney test).

### M1/M2 macrophage polarization in the MIA-induced OA model

Since the bi-directional links between local and systemic inflammation are poorly understood in OA, we examined the M1/M2 polarization in the spleen of MIA treated mice (Fig 2A and 3A). MIA treatment did not modify the percentage of M1 or M2 macrophages (F4/80^+^ CD11b^+^ iNOS^+^ CD38^+^ and F4/80^+^ CD11b^+^ CD206^+^ Arg1^+^, respectively) (Fig 2B and 3B). Similarly, we did not observe any change in the expression levels of the M1 and M2 specific markers. As previously observed in whole macrophages, a strong tendency towards downregulation of MHCII molecules and an upregulation of CD11b molecules were noted on M1 macrophages after MIA treatment (MHCII MFI Sham 5033 [4116-6280] *vs* MIA 2830 [2207-3804], *p=*0.0571; CD11b MFI Sham 14030 [11832-15270] *vs* MIA 30358 [16844-48337], *p=*0.0286) (Fig 2C). Interestingly, the frequency of M1 expressing MHCII molecules is reduced in mice with MIA-induced OA (Fig 2D).

**Figure 2.**
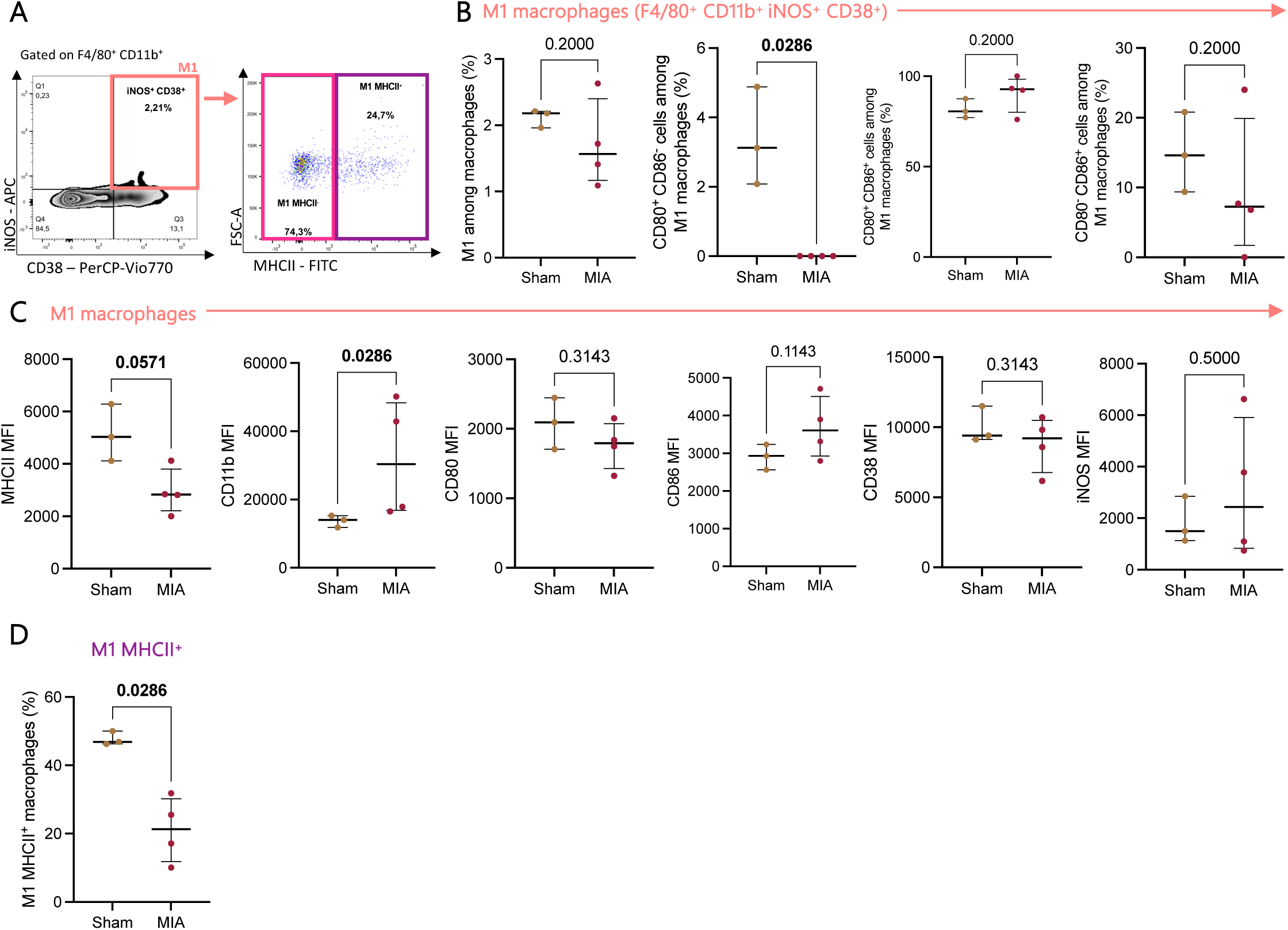
Reduction of MHCII⁺ M1 macrophage frequency and modulation of their activation profile in the MIA-induced OA model. (**A**) Representative gating strategy for M1 macrophages (F4/80⁺ CD11b⁺ iNOS⁺ CD38⁺) and their subsets according to MHCII expression (MHCII⁺ and MHCII^-^ M1 macrophages). (**B**) Quantification of total M1 macrophages among splenic macrophages (%) and analysis of activation markers (CD80, CD86) within this population. (**C**) Surface expression levels (MFI) of MHCII, CD11b, CD80, CD86, CD38, and iNOS in M1 macrophages. (**D**) Frequency of MHCII⁺ M1 macrophages in MIA mice and sham controls. Data are shown as individual values with median [25% percentile-75% percentile] (Sham n = 3; MIA n = 4; Mann-Whitney test).

**Figure 3.**
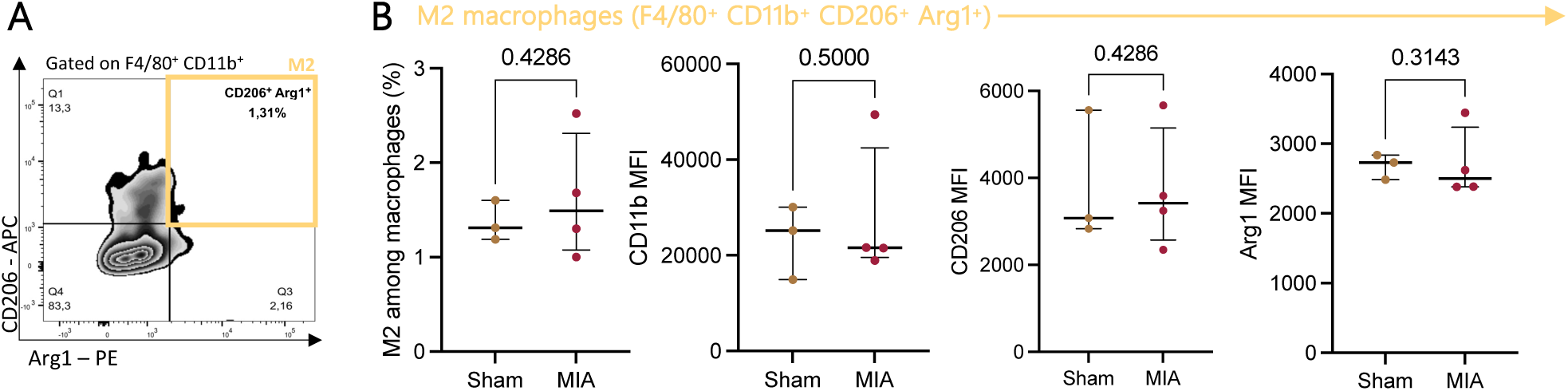
Splenic M2 macrophages are unaffected in the MIA-induced OA model. (**A**) Representative gating strategy for M2 macrophages (F4/80⁺ CD11b⁺ CD206⁺ Arg1⁺). (**B**) Quantification of total M2 macrophages among splenic macrophages (%) and analysis of surface marker expression levels (MFI) for CD11b, CD206, and Arg1. Data are shown as individual values with median [25% percentile-75% percentile] (Sham n = 3; MIA n = 4; Mann-Whitney test).

### Splenic DC distribution in the MIA-induced OA model

We performed phenotypic characterization of DCs in the OA condition, with a particular focus on conventional DC (cDCs) subsets (cDC1: F4/80^−^ CD11c^+^ CD11b^−^ MHCII^+^ and cDC2: F4/80^−^ CD11c^+^ CD11b^+^ MHCII^+^) (Fig 4A). The frequency of total DCs (F4/80^-^ CD11c^+^, including MHCII^+^ conventional and MHC^-^ tolerogenic DCs) was significantly increased in the MIA-treated group compared to sham controls (% total DCs Sham 5.80 [4.56-6.04] *vs* MIA 6.51 [6.22-6.89], *p=*0.0286) (Fig 4B). In MIA treated mice, we observed a decrease of cDC1 but not cDC2 frequencies (% cDC1 Sham 21.70 [19.10-25.20] *vs* MIA 11.05 [10.23-13.38], *p=*0.0286; % cDC2 Sham 1.17 [0.78-1.31] *vs* MIA 1.06 [0.77-1.16], *p=*0.2286) (Fig 4C, E). In MIA mice, expression of MHC class II molecules was also decreased in DCs, more marked in cDC2s than in cDC1s (MHCII MFI cDC2 Sham 16122 [14129-19827] *vs* MIA 4588 [4243-10868], *p=*0.0286; MHCII MFI cDC1 Sham 19452 [17896-19545] *vs* MIA 16415 [15528-18568], *p=*0.0571) (Fig 4D, F). CD11b molecules were upregulated in cDC2s while CD11c molecules were downregulated in cDC2s but not in cDC1s, in mice treated with MIA (CD11b MFI cDC2 Sham 14431 [11508-15091] *vs* MIA 58088 [54740-66600], *p=*0.0286; CD11c MFI cDC2 Sham 4534 [2913-5630] *vs* MIA 2103 [1722-2336], *p=*0.0286; CD11c MFI cDC1 Sham 4781 [4582-4802] *vs* MIA 4209 [3941-5216], *p=*0.2000) (Fig 4D, F). The cDC2 but not cDC1 had a higher frequency of the CD80^-^ CD86^+^ subset in MIA mice (Fig 4C, E).

**Figure 4.**
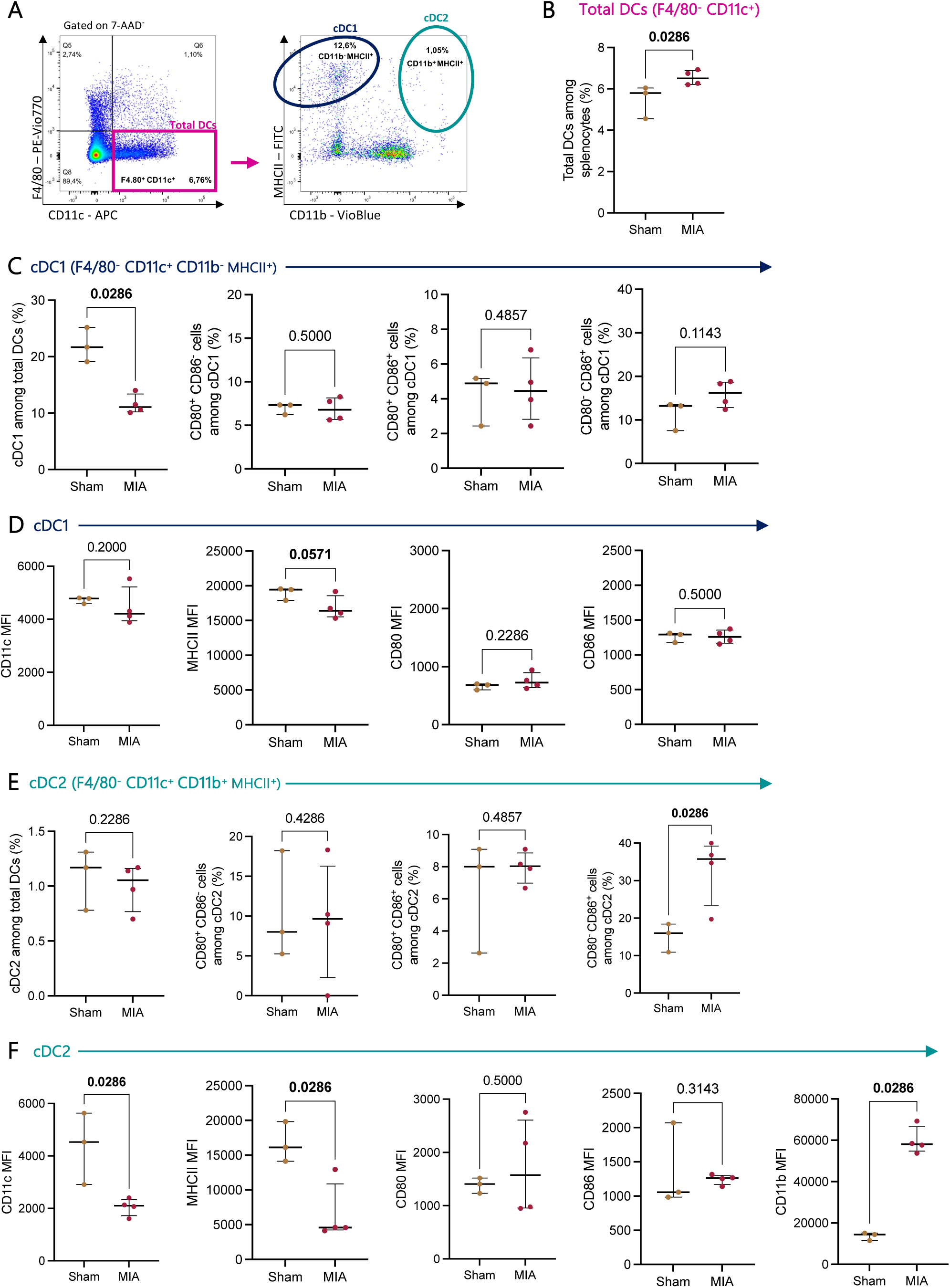
Modulation of splenic dendritic cell subsets and their activation profiles in the MIA-induced OA model. (**A**) Representative gating strategy for total dendritic cells (DCs; F4/80⁻ CD11c⁺) and their subsets: conventional DC subsets: cDC1 (F4/80⁻ CD11c⁺ CD11b⁻ MHCII⁺) and cDC2 (F4/80⁻ CD11c⁺ CD11b⁺ MHCII⁺). (**B**) Quantification of total DCs among splenocytes (%) in MIA mice and sham controls. (**C-D**) Characterization of cDC1. Quantification of total cDC1, CD80^+^ CD86^+^ and CD80^-^ CD86^+^ cDC1 among splenocytes (%) and analysis of surface marker expression (MFI - CD11c, MHCII, CD80 and CD86). (**E-F**) Quantification of total cDC2, CD80^+^ CD86^+^ and CD80^-^CD86^+^ cDC1 among splenocytes (%) and analysis of surface marker expression (MFI - CD11c, CD11b, MHCII, CD80 and CD86). Data are shown as individual values with median [25% percentile-75% percentile] (Sham n=3; MIA n=4; Mann-Whitney test).

Tolerogenic DCs that did not express MHCII molecules were subdivided into three subsets according to CD11b expression level (Fig 5A). In MIA condition, the frequency of CD11b^-^ MHCII^-^ DCs was decreased whereas the percentage of CD11b^low^ MHCII^-^ DCs was increased (% CD11b^-^ MHCII^-^ DCs Sham 30.40 [29.50-31.80] *vs* MIA 21.35 [19.35-22.98], *p=*0.0286; % CD11b^low^ MHCII^-^ DCs Sham 42.10 [41.10-47.20] *vs* MIA 63.50 [60.30-63.78], *p=*0.0286) (Fig 5B, D). In addition, CD11b^-^ MHCII^-^ DCs had a lower level of CD86 molecules, while CD11b^low^ MHCII^-^ DCs showed a slight tendency toward increased CD86 expression and had a higher level of CD11b molecules (CD86 MFI CD11b^-^ MHCII^-^ Sham 1516 [1501-1581] *vs* MIA 1354 [1296-1444], *p=*0.0286; CD86 MFI CD11b^low^ MHCII^-^ Sham 761 [733-766] *vs* MIA 804 [773-809], *p=*0.0571; CD11b MFI CD11b^low^ MHCII^-^ Sham 3634 [3626-3818] *vs* MIA 4274 [4042-4620], *p=*0.0286) (Fig 5C, E). CD11b^+^ MHCII^-^ DCs had a significantly higher proportion of co-expressing CD80^+^ CD86^+^ in MIA treated mice (% CD11b^+^ MHCII^-^ CD80^+^ CD86^+^ Sham 0.00 [0.00-3.23] *vs* MIA 10.80 [5.90-18.05], *p=*0.0286) (Fig 5F). As observed for cDC2s in MIA condition, CD11b^+^ MHCII^-^ DCs expressed a lower level of CD11c and higher level of CD11b (CD11c MFI CD11b^+^ MHCII^-^ Sham 1917 [1824-2414] *vs* MIA 1506 [1366-1595], *p=*0.0286; CD11b MFI CD11b^+^ MHCII^-^ Sham 41518 [37830-46139] *vs* MIA 68637 [65063-70315], *p=*0.0286) (Fig 5G).

**Figure 5.**
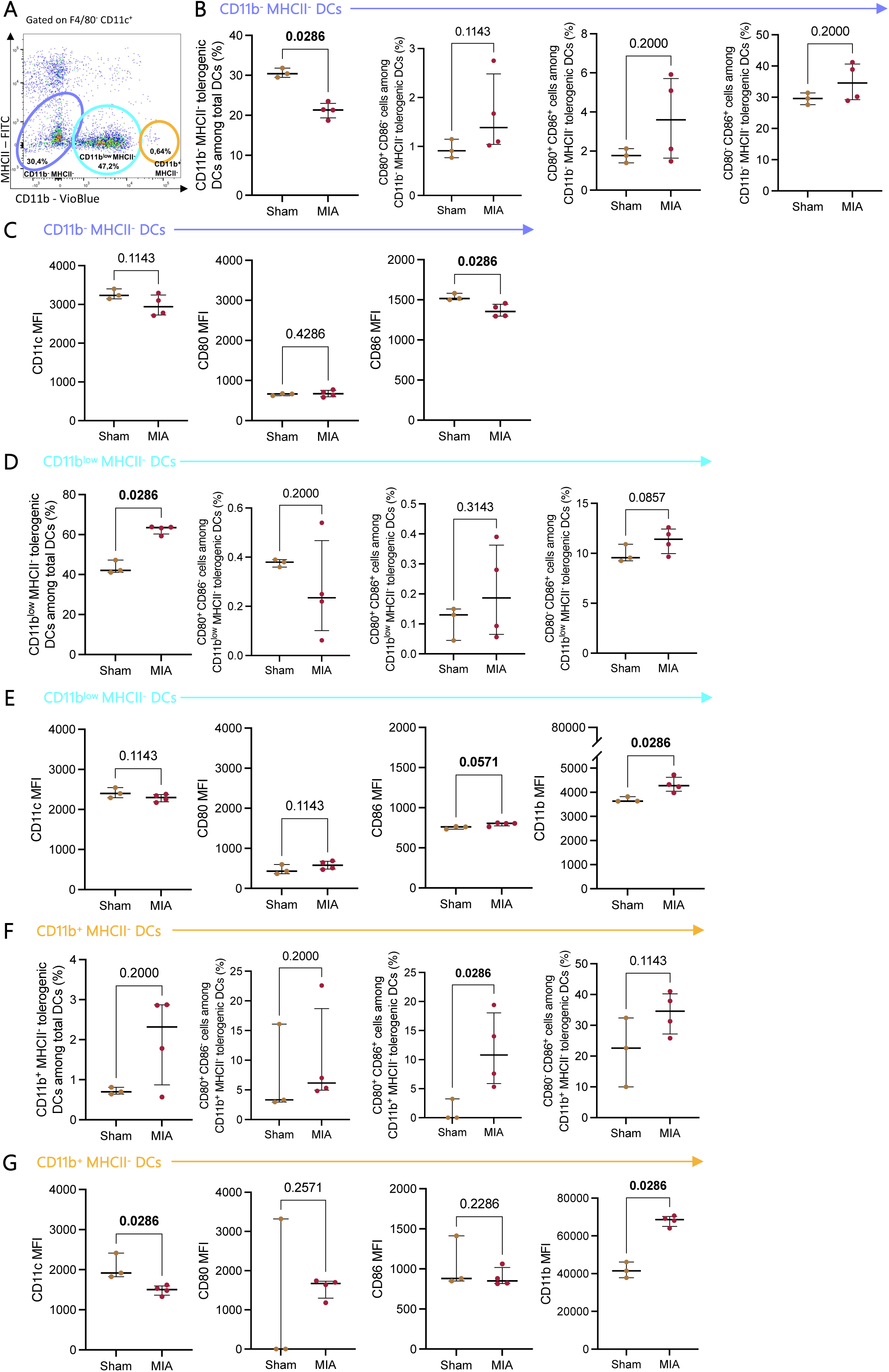
Modulation of splenic tolerogenic dendritic cell subsets in the MIA-induced OA model. (**A**) Representative gating strategy for tolerogenic dendritic cells (F4/80^-^ CD11c⁺ MHCII^-^), and their subsets according to CD11b expression into CD11b^-^ MHCII^-^, CD11b^low^ MHCII^-^, and CD11b^+^ MHCII^-^ DCs. (**B-C**) Quantification of total CD11b^-^MHCII^-^ DCs, CD80^-^ CD86^+^, CD80^+^ CD86^+^ and CD80^-^ CD86^+^ CD11b^-^ MHCII^-^ DCs among splenocytes (%) and analysis of surface marker expression (MFI - CD11c, CD80 and CD86). (**D-E**) Quantification of total CD11b^low^ MHCII^-^ DCs, CD80^-^ CD86^+^, CD80^+^ CD86^+^ and CD80^-^ CD86^+^ CD11b^low^ MHCII^-^ DCs among splenocytes (%) and analysis of surface marker expression (MFI - CD11b, CD11c, CD80 and CD86). (**F-G**) Quantification of total CD11b^+^ MHCII^-^ DCs, CD80^-^ CD86^+^, CD80^+^ CD86^+^ and CD80^-^ CD86^+^ CD11b+ MHCII^-^ DCs among splenocytes (%) and analysis of surface marker expression (MFI - CD11b, CD11c, CD80 and CD86). Data are shown as individual values with median [25% percentile-75% percentile] (Sham n = 3; MIA n = 4; Mann-Whitney test).

### T and B cell distribution in the MIA-induced OA model

Finally, we performed phenotypic characterization of lymphoid cells, including leukocytes, B and T cells (Fig 6A, D). The proportion of leukocytes and B cells was decreased in MIA mice (% CD45 Sham 98.60 [98.40-98.60] *vs* MIA 81.60 [50.23-88.38], *p=*0.086; % B cells Sham 82.00 [80.50-84.00] vs MIA 57.70 [28.93-64.28], p=0.0286) (Fig 6B, C) whereas no difference was observed for T cells (% T cells Sham 21.90 [21.50-27.50] *vs* MIA 28.75 [22.38-29.50], *p=*0.1714) (Fig 6E). In the same mice, we showed a decrease of CD4^+^ T cells and CD4^+^ regulatory T cells (Tregs); and conversely an increase of CD8^+^ T cells (%CD4^+^ T cells Sham 66.00 [63.70-68.20] *vs* MIA 57.05 [52.68-59.40], *p=*0.0286; % CD4^+^ Tregs Sham 0.71 [0.56-0.95] *vs* MIA 0.09 [0.02-0.22], *p=*0.0286; % CD8^+^ T cells Sham 22.00 [21.50-22.90] *vs* MIA 32.85 [31.78-35.58], *p=*0.0286) (Fig 6F, G). Additionally, CD4^+^ T cells displayed a lower level of the activation markers CD25, CD44 and CD69 in MIA condition (% CD4^+^ CD25^+^ T cells Sham 11.60 [11.40-12.30] *vs* MIA 6.83 [6.41-7.52], *p=*0.0286; % CD4^+^ CD44^+^ T cells Sham 22.30 [22.00-25.00] *vs* MIA 15.25 [15.03-15.63], *p=*0.0286; % CD4^+^ CD69^+^ T cells Sham 10.50 [9.74-11.40] *vs* MIA 8.11 [7.97-8.81], *p=*0.0286) (Fig 6F). For CD8^+^ T cells, only the activation marker CD25 was downregulated in MIA mice (% CD8^+^ CD25^+^ T cells Sham 4.24 [3.23-4.41] *vs* MIA 2.34 [1.98-2.88], *p=*0.0286) (Fig 6G).

**Figure 6.**
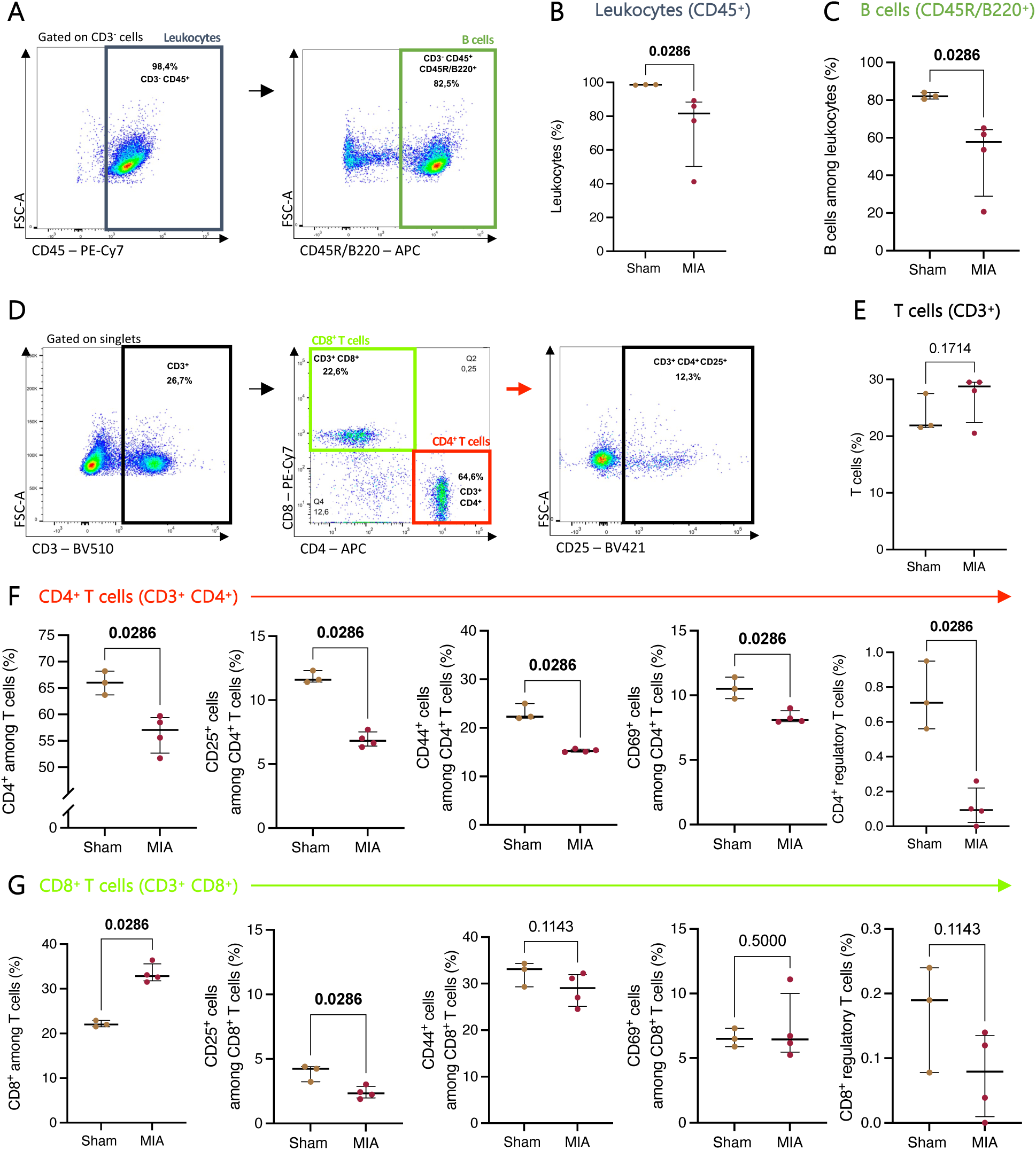
Reshaping of the splenic adaptive immune compartments toward reduced B and T cell activation in the MIA-induced OA model. (**A**) Representative gating strategy for leukocytes (CD3^-^ CD45⁺) and B cells (CD3^-^CD45⁺ CD45R/B220⁺) among splenocytes. (**B-C**) Quantification of total leukocytes and B cells in MIA mice and sham controls. (**D**) Representative gating strategy for T cells (CD3⁺) and their subsets (CD4⁺ and CD8⁺). (**E**) Total T cell frequency. (**F**) Quantification of CD4⁺ T cells (CD3⁺ CD4⁺) and their activation markers CD25⁺, CD44⁺and CD69⁺. Quantification of CD4⁺ regulatory T cells (CD4^+^ CD25⁺ FoxP3⁺). (**G**) Quantification of CD8⁺ T cells (CD3⁺ CD4⁺) and their activation markers CD25⁺, CD44⁺, and CD69⁺. Quantification of CD8⁺ regulatory T cells (CD8^+^ CD25⁺ FoxP3⁺). Data are shown as individual values with median [25% percentile-75% percentile] (Sham n = 3; MIA n = 4; Mann-Whitney test).

## Discussion

Interactions between local joint inflammation and the systemic immune response is poorly understood in chronic OA. Altogether, using a detailed immunophenotyping of splenocytes, our data reveal a remodeling and plasticity of the systemic innate and adaptative immune response in chronic OA.

Macrophages are central players in OA as active regulators of inflammation, tissue remodeling, and pain. In the MIA-induced OA model, splenic macrophages (including M1 macrophages) and also cDC1 and cDC2 displayed a marked downregulation of MHCII expression level. Interestingly, we reported the same effect in an acute inflammation mouse model triggered by *in vivo* injection of lipopolysaccharide (LPS) [25]. Given the function of MHCII molecules in Ag presentation, this might reflect a negative feedback mechanism to prevent chronic T cell activation in OA. However, a previous study described an increase of MHCII molecules (HLA-DR) on circulating monocytes in human OA [20]. This apparent discrepancy likely reflects species-specific immune regulation, as well as differences in disease stage and systemic environment. In mice, macrophages may transiently downregulate MHCII as part of a resolution program in OA, whereas in human late-stage OA joints, chronic pro-inflammatory signaling maintains or increases MHCII expression.

In a study based on genome wide data sets analyzed by deconvolution, Chen et al. (2020) describe increased proportions of “activated” dendritic cells in OA synovial tissue, suggesting a shift in immune cell activation status [11]. However, the criteria used to define activation versus resting states are not explicitly detailed, making it challenging to directly compare findings across studies. Here, we used flow cytometry that is a robust method to quantify rare immune cell subsets measuring expression of cell surface markers. We distinguished conventional DCs (cDCs) based on MHCII and CD11b expression, allowing separation into cDC1 (CD11b⁺ MHCII⁺) and cDC2 (CD11b^-^ MHCII⁺) subsets. In addition, we identified a major population (∼80% of total DCs) lacking MHCII expression (F4/80^-^ CD11c⁺ MHCII^-^), which we defined as “tolerogenic DCs” based on their immature phenotype. Interestingly, we further resolved these tolerogenic DCs into three subsets based on CD11b expression: CD11b^-^, CD11b^low^, and CD11b^+^. In OA animals, we observed an overall increase in total DCs (including conventional and tolerogenic DCs). However, this expansion does not appear to result from an increase in mature conventional DCs, as cDC1 were significantly decreased and cDC2 remained unchanged. Instead, the increase in total DCs seems primarily driven by changes within the tolerogenic DC compartment. Notably, while the proportion of CD11b^-^ tolerogenic DCs was reduced, CD11b^low^ tolerogenic DCs markedly expanded in OA animals, rising from approximately 40% in controls to over 60% in the OA group. The frequency of CD11b^+^ tolerogenic DCs, in contrast, remained unchanged between groups.

Importantly, these frequency changes were mirrored by modulation of activation markers. The expression of CD86 level on tolerogenic DC subsets closely followed the abundance trends of their respective populations: decreased in CD11b^-^ DCs, increased in CD11b^low^ DCs, and unchanged in CD11b^+^ DCs. We also noticed an upregulation of this costimulatory molecule on whole macrophages. These observations suggest a coordinated regulation of both abundance and immunological function. In addition to being an activation marker, CD86 has been shown to play a central part in sustaining prolonged interactions between antigen-presenting cells and T cells [26]. The modulation of CD11b-defined tolerogenic DC subsets and their CD86 expression suggests functional plasticity within this compartment. The increase in CD11b^low^ cells, displaying higher CD86 levels, may reflect an intermediate activation state with preserved immunoregulatory capacity, possibly induced by the chronic, low-grade inflammation characteristic of OA. Such phenotypic flexibility is a hallmark of DC biology, especially in tissues requiring long-term immune homeostasis [27].

Interestingly, CD11b, also known as integrin αM, is traditionally recognized as part of the CD11b/CD18 heterodimer (also called Mac-1), which plays a central role in cell adhesion, migration, and phagocytosis, particularly within the innate immune system [28]. However, more recent studies have uncovered a distinct role for CD11b in immune modulation. Notably, CD11b has been shown to act as a functional co-receptor with annexin A2 for tissue plasminogen activator (tPA), promoting NF-κB-dependent macrophage activation in models of chronic kidney inflammation [29,30]. We also explored the systemic adaptive immune landscape in OA by characterizing B and T cell subsets in the spleen. Total leukocyte frequency (CD45^+^) was significantly reduced in OA animals, a decrease that appeared to be primarily driven by a marked reduction in B cells (CD45R^+^/B220^+^), which dropped from approximately 80% to less than 60% of total splenocytes. In contrast, total T cell (CD3^+^) frequencies were unaffected. However, further subset analysis revealed a significant shift in the CD4/CD8 T cell balance: we observed a decrease in CD4^+^ T cells, including activated subsets (CD69^+^, CD44^+^, CD25^+^), and a reduction in regulatory T cells (CD4^+^ CD25^+^ Foxp3^+^). Conversely, CD8^+^ T cells were increased in OA animals, with a selective rise in CD25^+^ CD8^+^ T cells, indicative of sustained or late activation. These changes highlight a differential impact of OA on CD4^+^ and CD8^+^ compartments and suggest an altered systemic immune tone, skewed toward cytotoxic over helper/regulatory functions.

The reduction in total leukocytes and particularly B cells within the spleen may reflect chronic immune exhaustion or redistribution of lymphocyte subsets toward local inflammatory sites such as the joint, draining lymph nodes, or bone marrow. While B cells are considered relatively minor players in the synovial compartment, representing only ∼5% of the total infiltrate in OA joints [19], they remain a dominant population in secondary lymphoid organs like the spleen. Here, their reduction could signal impaired B cell maintenance or an OA-induced systemic contraction of the humoral compartment. Although our gating strategy using CD45R/B220 does not allow the distinction of the maturation stages, the drop in global B cell frequency raises the possibility of altered B cell homeostasis or survival in response to chronic inflammation.

On the T cell side, the observed decrease in CD4^+^ T cells and their activated and regulatory subsets, combined with the expansion of activated CD8^+^ T cells, points to a shift toward a cytotoxic systemic immune profile in OA. This imbalance may reflect disrupted immune homeostasis and a loss of regulatory control, potentially exacerbating chronic low-grade inflammation. Importantly, by integrating these observations with our phenotypic profiling of myeloid cells, our study underscores a broader systemic immune remodeling in OA. It also opens avenues for exploring the functional crosstalk between innate (DCs/macrophages) and adaptive (T/B cells) compartments in shaping chronic joint inflammation.

Taken together, the remodeling observed in both the myeloid and lymphoid compartments suggests that chronic OA may drive a coordinated reprogramming of systemic immune responses. The marked modulation of tolerogenic DC subsets, characterized by frequency shifts in parallel with changes in surface CD86 expression, highlights a dynamic adjustment of their immunoregulatory potential. This is accompanied by reduced MHCII expression on conventional DCs and macrophages, suggesting a broader alteration of antigen presentation capacity. In parallel, the decline in CD4^+^ T cells, including activated and regulatory subsets, and B cells, along with the selective increase in activated CD8^+^ T cells, points to a systemic polarization toward cytotoxic responses. These patterns raise the hypothesis that DC plasticity in OA not only reflects local adaptation to inflammatory cues but may actively influence adaptive immune fate through modified co-stimulatory signaling, cytokine production, or impaired tolerogenic instruction. Altogether, our data point to a functionally interconnected remodeling of innate and adaptive immunity, where altered DC phenotypes could act as central coordinators of immune imbalance in chronic OA.

## Declaration

### Ethics approval

Animal experiments were conducted in strict accordance with European Directive 2010/63/EU and French regulations (Decree 87/848) at the CURB facility (Centre Universitaire de Ressources Biologiques, accreditation #D14118001). All procedures received prior authorization from the French Ministry of Education and Research as well as approval from the regional ethics committee CENOMEXA (APAFIS authorization #16185). Experimental procedures complied with the ARRIVE guidelines (Animal Research: Reporting of In Vivo Experiments; https://www.nc3rs.org.uk).

### Consent for publication

Not applicable

### Availability of data and materials

The datasets supporting the conclusions of this article are included within the article.

### Authors’ contributions

Conceptualization: C.B. and O.T.; Methodology: C.B., O.T., C.S., S.B., B.LM.; Validation: C.B., O.T., C.S., S.B.; Formal analysis: C.S.; Investigation: C.S., and S.B.; Resources: C.B. and O.T.; Data Curation: C.S.; Writing - Original Draft: C.S., C.B and O.T.; Writing - Review & Editing: All authors; Visualization: C.S.; Supervision: C.B., O.T., K.B., D.V., and B.LM.; Project administration: C.B. and O.T.; Funding acquisition: C.B., O.T., K.B., D.V., and B.LM.

### Declaration of generative AI and AI-assisted technologies in the writing process

During the preparation of this work the authors used ChatGPT in order to improve English editing of some parts of the manuscript. After using this tool/service, the authors reviewed and edited the content as needed and take full responsibility for the content of the published article.

## Acknowledgments

We thank Véronique Agin (PhIND, Caen), Palma Pro and all animal facility staff (CURB, Unicaen, France) for technical assistance.

